# Telomere length across land plants tracks warm season temperature and not life history

**DOI:** 10.64898/2026.09.18.752638

**Authors:** Linh Nguyen, Jae Young Choi

## Abstract

Telomeres protect chromosome ends, yet their length varies widely among organisms. What shapes this variation is a long-standing question with implications for chromosome function and evolution. In animals, telomere length variation is explained by life-history differences related to senescence and tumorigenesis but the drivers in plants are unknown. Because plants are sessile organisms where growth depends directly on their surroundings the telomere length might be under selection from environmental factors. We tested this alternative ecological hypothesis by estimating telomere lengths from long read sequencing data for 264 species spanning the green plant kingdom. Telomere length differed among major lineages where green algae had the shortest telomeres and seed plants the longest. But plants with different life histories differed little in telomere length. Instead, after accounting for phylogeny and sequencing read length, telomere length increased with the peak temperature of the warmest month by 16% per 10 °C. Peak temperature explained telomere length better than latitude or other climatic variables. This covariation was largely phylogenetically driven and present in both recently and anciently diverged lineages. Domestication, which was proposed to lengthen telomeres, had no consistent effects when we compared crops with their wild progenitors. We conclude in contrast to animal telomeres, the life-history traits we analyzed in this study explained little of the variation in plant telomere length. Alternatively, we propose the ecological adaptation hypothesis where telomere length in plants is shaped by selection relating to the thermal environment of growth.

## Introduction

Telomeres are nucleoprotein structures composed of repetitive DNA and specialized proteins that are found at chromosome ends. Telomeres have the crucial role of protecting chromosome ends from degradation and instability (O’Sullivan and Karlseder 2010; Stewart et al. 2012).

During DNA replication, the DNA polymerase cannot properly replicate the ends of the lagging strand DNA, and without any intervention results in the loss of genetic material after each round of cellular division (*i.e.* the end replication problem (Watson 1972; Olovnikov 1973)). In addition, exposed chromosome ends can trigger the DNA damage response pathway, resulting in cell cycle arrest and disrupting chromosome integrity (de Lange 2018). To prevent such problems, the telomerase maintains the chromosome ends and specialized telomere binding proteins shelter and protect the chromosome ends (Shay and Wright 2019). Maintaining an optimal telomere length is essential for genome stability, as abnormally long telomeres are associated with cancer cells and tumorigenesis (Savage 2024), whereas shortened telomeres are associated with aging and adverse effects on lifespan (Armanios 2009). Because of the detrimental effects arising from changes in the telomere length, it is predicted that length of the telomere should be maintained within an optimal range and evolutionary conservation should prevent the variation of telomere lengths between organisms.

Yet, despite the highly deleterious consequences of telomere shortening and lengthening telomere length is not conserved. For example, telomere length can vary over two fold between two human individuals (Demanelis et al. 2020; Karimian et al. 2024), and across the tree of life even greater levels of variation is observed between species, like yeast (*Saccharomyces cerevisiae*) with a telomere length of 360 bp (Garrido et al. 2026) and mouse (*Mus musculus*) with telomere lengths of over 20 kbp (Zijlmans et al. 1997). Why telomere length varies among natural organisms remains a largely open question. Quantitative genetics studies in both plant and animals have shown telomere length differences between individuals are highly heritable (Zhu et al. 1998; Gatbonton et al. 2006; Codd et al. 2013; Abdulkina et al. 2019), suggesting telomere length has great potential to be shaped by evolutionary change through natural selection (Lynch and Walsh 1998). Identifying the selective force has long interested evolutionary and chromosome biologists (Monaghan et al. 2018), as it is key to understanding why telomere length varies in nature.

Cross-species analysis has been a crucial approach for our understanding of the potential driver of selection on telomere length variation. Many of these studies have focused on vertebrates, hence the findings cannot be generalized for all organisms, nevertheless these studies found association between telomere length and various life-history traits such as body size, life span, and metabolic strategy among species (Heidinger et al. 2012; Olsson, Wapstra, and C. Friesen 2018; Pepke and Eisenberg 2022; Benson et al. 2025). Additional studies have found activity of the telomerase or the attrition rate of the telomere is also associated with life-history traits (Haussmann et al. 2003; Gomes et al. 2011; Dantzer and Fletcher 2015; Sudyka et al. 2016; Tricola et al. 2018; Whittemore et al. 2019; Pepke et al. 2020; Criscuolo et al. 2021). A major conclusion from these studies is that telomere length shortens with increasing age and predicts aspects of aging and lifespan. Telomere length is also associated with reproductive success, fitness, and stress response (Epel et al. 2004; Haussmann et al. 2005; Bauch et al. 2014; Asghar et al. 2015), suggesting telomeres might have an underappreciated role in the evolution of life-history traits. Studying telomere dynamics in the context of ecology and evolution (Monaghan and Haussmann 2006; Monaghan 2010; Gomes et al. 2011; Monaghan 2014; Olsson, Wapstra, and C.R. Friesen 2018; Sudyka 2019) is crucial for gaining a deeper understanding of the factors that maintain telomere length in natural populations.

Several hypotheses have been proposed to explain the associations between telomeres and life-history or fitness-related traits (Tobler et al. 2022), which can be grouped into two major models. First, the tumor suppression model hypothesizes replicative senescence, the cellular senescence arising from shortening of the telomere, is an adaptive mechanism to prevent the formation of precancerous cells (Campisi 2001). Organisms with large bodies have an increased number of cells, which can increase the opportunity of developing cancerous cells. In addition, organisms with long lifespans will have an increased time period of accumulating mutations associated with tumorigenesis (Artandi and DePinho 2010; Aviv et al. 2017). Here, the variation in telomere length among organisms arose as an adaptive consequence of an anti-cancer strategy (de Lange and Jacks 1999; Shay 2016). On the other hand, telomere length variation could be an evolutionary consequence from the varying life-history strategies.

Organisms have evolved differing schedules of growth, reproduction, and survival (*i.e.* life-history strategy) that are most optimal to its surrounding ecology (Metcalf and Pavard 2007). Resources are not infinite in nature and trade-offs constrain the optimization of trait combinations for an organism, resulting in the diversity of life-history strategies across the tree of life (Stearns 1998). The pace of life model proposes telomere length variation arises as a consequence of the differing life-history strategies (Giraudeau et al. 2019). Organisms on the fast pace of life axis grow rapidly, reproduce early, and die young, whereas on the opposite side of the axis organisms grow slowly, reproduce late, and die older (Bielby et al. 2007; Salguero-Gómez et al. 2016; Healy et al. 2019). Under a fast pace of life strategy, resources are concentrated on growth and reproduction but at the cost of self maintenance leading to an early onset of senescence (Giaimo and Traulsen 2019; Cayuela et al. 2020). Increased reproductive effort is associated with shortened telomeres in multiple species (Kotrschal et al. 2007; Bauch et al. 2013; Sudyka et al. 2014; Sudyka et al. 2019; Morland et al. 2023), suggesting trade-offs in life-history traits can result in the variation of telomere length. The shortened length of the telomere could be the direct cause of senescence or telomere length may be a physiological marker of an organism and the association is simply indirect (Young 2018).

Animal studies were instrumental in establishing the foundational framework for understanding the evolutionary processes that shape telomere length in natural populations. In plants, however, far less is known on what selective force is shaping their telomeres. While components of the telomere and the telomere-maintenance complex are largely conserved between plants and animals (Watson and Riha 2010), it is unclear if telomere length variation in both kingdoms are driven by the same ecological and evolutionary mechanism. In plants, for instance, because of the presence of a cell wall cancerous cells are predicted to have far less deleterious consequences (Doonan and Sablowski 2010). Hence, telomere length variation in plants is unlikely to be driven by the tumor suppression model. On the other hand, shared life-history trade-offs can cause similar life-history strategies even between deeply diverged species (Stott et al. 2024). The pace of life model could also drive telomere length in plants, although the eco-evolutionary driver of selection is likely to differ between plants and animals. One major difference is rooted in how organismal development arises from the initial stem cell. Unlike animals where stem cell based development is confined to specific tissues or early developmental stages, growth of plants are indeterminate and the entire body plan is generated throughout its lifespan through a population of stem cells called the meristem (Weigel and Jürgens 2002). The protein complexes maintaining the telomere are primarily active in the meristem (Fitzgerald et al. 1996), suggesting telomere maintenance could be coupled to meristem maintenance that ultimately influences the plant body plan development. Past empirical studies have discovered telomere length is correlated with flowering time in *Arabidopsis*, maize, and rice (Choi et al. 2021), meanwhile *A. thaliana* mutants with genetically manipulated telomere lengths had changes in flowering time and reproductive output during stress (Campitelli et al. 2022). These studies suggest a potential biological link between the telomere and life-history traits, indicating telomere length is an understudied trait associated with the evolution of life-history strategy in plants. For instance, perennials that undergo multiple rounds of vegetative and reproductive growth, meristem maintenance is essential (Albani and Coupland 2010) and telomere maintenance could have had an equally important role during the evolution of a perennial life-history.

An alternative hypothesis proposes plant telomere length is shaped by the environment. Plants are sessile and the rates of cell division, meristem growth, and overall organismal development depends strongly on the ambient environment (Bennett et al. 1982; Francis and Barlow 1988). Other fundamental chromosomal properties of plants already track climatic variation. Genome size, for example, varies with temperature across species (Bureš et al. 2024), meanwhile telomere length itself shows hints of correlation with the environmental variation in plants. For example, telomere length in pine varies with seasonal temperature (Mu et al. 2015) and *A. thaliana* ecotypes from lower latitudes have longer telomeres (Choi et al. 2021). Under the ecological hypothesis, we predict telomere length should vary with the climate and particularly with the environmental parameters that are crucial during growth.

In this study we tested the life-history and ecological hypotheses for telomere length variation in plants using a comparative biology approach. We generated the largest telomere length dataset to date, encompassing 264 species spanning the entire green plant kingdom. We then detailed each species molecular traits (diploidy vs. polyploidy, genome size, and chromosome number), life-history traits (annual vs. perennial, herbaceous vs. woody, height, and body plan sizes), and ecology (altitude, latitude, bioclimatic variables, and range size), and tested its association with telomere length using phylogenetic comparative methods (Cornwallis and Griffin 2024) and multi-response phylogenetic mixed models (Halliwell et al. 2025). As a further test of the life-history hypothesis, we compared telomere lengths of domesticated crops to their wild progenitors, as past studies have suggested selection on the life-history variation relating to domestication has lengthened the telomeres of domesticated organisms (Pepke and Eisenberg 2022; Colt et al. 2024). Our results show plant telomeres contrast with animal telomeres, where life-history differences and domestication did not explain the length variation, indicating a fundamentally different evolutionary force was shaping telomere lengths in plants. Instead, we discovered that temperature, particularly the peak temperature of the warmest month was associated with global plant telomere length variation. Importantly, temperature explained telomere length better than latitude or other climatic variables. We propose selection arising from the thermal environment of growth, rather than life history, shapes telomere length variation across the plant kingdom.

## Results

### Plant kingdom wide telomere length variation

Measuring telomere length using traditional techniques is highly laborious and low throughput, instead we estimated each plant species telomere length using a computational approach. Recently we developed Topsicle (Nguyen and Choi 2025), a novel computational method that analyzes long read sequencing data and searches for reads spanning both telomere and subtelomere regions to infer the telomere length. Topsicle was benchmarked against the terminal restriction fragment assay (Harley et al. 1990), a gold standard for measuring telomere length and had high correlations (r = 0.74). We took advantage of the abundant whole genome long read sequencing data for plants (Xie et al. 2024) and estimated telomere lengths for 264 species by analyzing a total of 692 sequencing libraries that comprised of 10 TB of data, 1.5 billion sequencing reads, and 18 trillion nucleotides (Table S1).

Because telomere length is inferred from long reads that span the telomere, the technical differences between sequencing libraries could bias the estimates. Contig N50 of the assembled genomes had no significant correlation with telomere length (Spearman ρ = −0.030, p = 0.66, n = 217; Table S2). But the average read (spot) length of each species’ SRA libraries was positively associated with telomere length after accounting for phylogeny (PGLS slope = 0.197 and p = 3.6e-5; Fig S1A and Table S2). Telomere lengths estimated from the Oxford Nanopore (ONT; 65 species) platform were also on average 1.28 fold longer than those from PacBio (192 species) platform (PGLS p = 1.3e-5; Fig S1B). Spot length and platform were each independently associated with telomere length (PGLS spot length p = 8.8e-4 and platform p = 3.0e-4). We therefore included log10 spot length as a covariate in all subsequent comparative analyses and tested the robustness of the results to sequencing platform.

### Telomere length differs among major plant lineages

Using the telomere length estimates from 264 plants we first investigated the deep evolutionary changes in the telomere length, by grouping the species into 6 clades and the spermatophytes (*i.e.* seed plant) were further divided into 16 orders (Fig 1A). Across the plant kingdom the median telomere length was 5,857.5 bp. We noticed telomere length increased with major evolutionary transitions towards terrestrialization. Specifically, land plants had significantly longer telomeres compared to green algae (median telomere length 6,024 bp vs. 590 bp and Mann Whitney U test p-value = 7.7e-6). Among land plants, vascular plants had significantly longer telomeres compared to bryophytes (median telomere length 6,102 bp vs. 1,710 bp and Mann Whitney U test p-value = 1.4e-4), and seed plants had significantly longer telomeres compared to bryophytes and the lycophyte (median telomere length 6,120 bp vs. 1,745 bp and Mann Whitney U test p-value = 5.0e-05).

**Figure 1.**
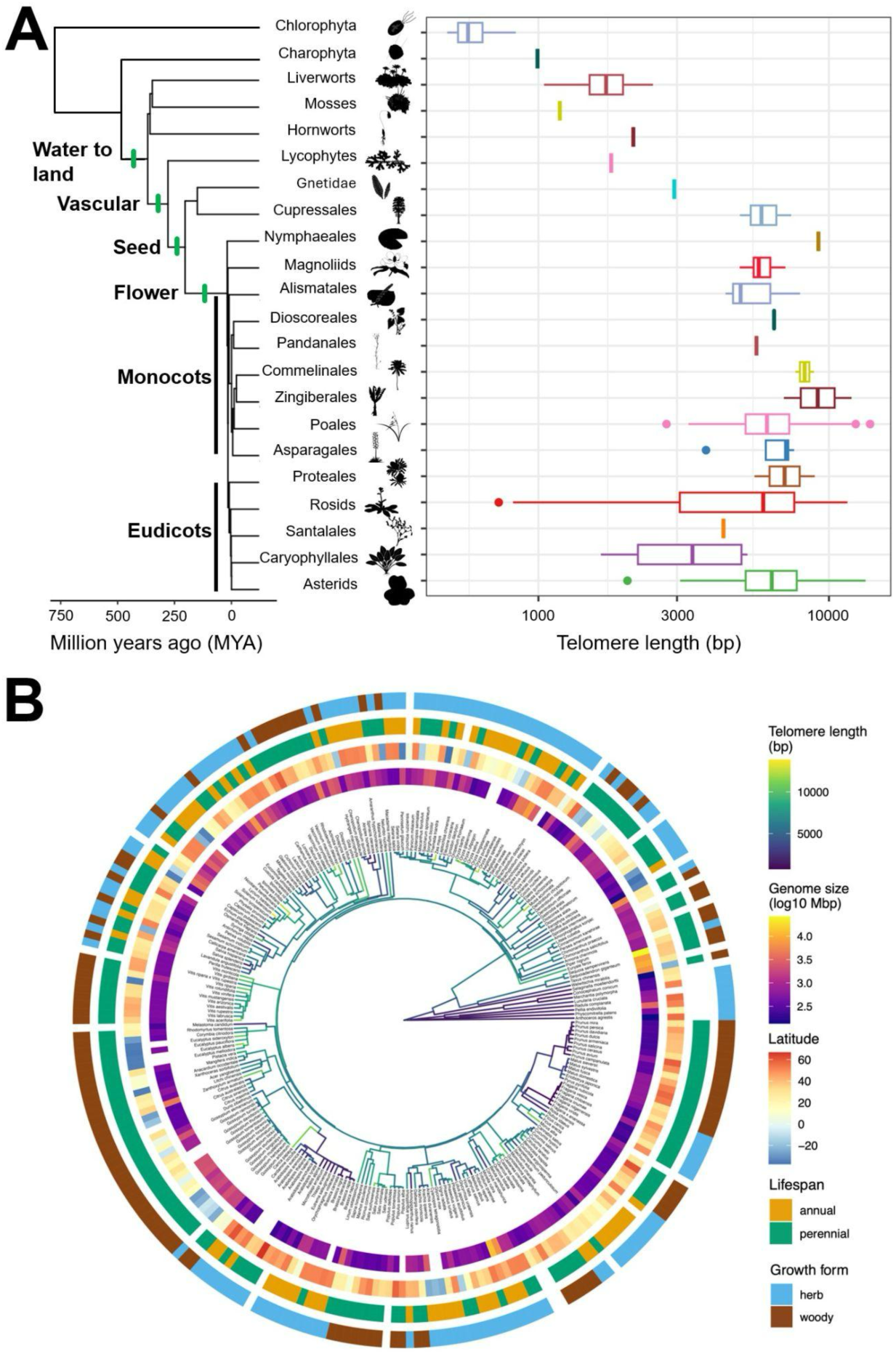
Telomere length variation across the plant kingdom. (A) Telomere length distribution for the major clades or orders across the plant kingdom. Silhouette images were obtained from PhyloPic. (B) Circular phylogenetic tree showing telomere length variation for the 257 land plant species. Outer rings display genome size, latitude, lifespan, and growth form.

Focusing on angiosperms, which comprised the largest sample size in our dataset (N = 245), monocots did not have significantly longer telomeres compared to eudicots (median telomere length 6,435 vs. 6,028 bp and Mann Whitney U test p-value = 0.072). Because each of these contrasts reflects a single major evolutionary transition, we interpreted them as descriptive patterns rather than as tests of the effect of the transition on telomere length. We therefore used phylogenetic comparative methods across all land plants to test the factors associated with telomere length variation.

### Phylogenetic signal in plant telomere lengt

We focused our analysis on embryophytes (*i.e.* land plant) to gain a deeper understanding of telomere length evolution in plants (Fig 1B) (N = 257). Initially, we determined the best fitting macroevolutionary model (Table S3) for telomere length and discovered the lambda model had the best fit (ΔAICc = 17.8 compared to second best fit model). The estimated phylogenetic signal (λ) was 0.90 (95% CI 0.84 – 0.94) and it was significantly different from 0 and 1. A λ value of 0 indicates no phylogenetic signal, while value of 1 indicates the trait is evolving under a pure Brownian motion model (Pagel 1997; Pagel 1999; Pearse et al. 2025), suggesting telomere length in plants has a strong phylogenetic signal but the trait was not evolving under a complete Brownian motion model and other evolutionary forces were involved.

Plants were then grouped by life-history strategy and we re-estimated the phylogenetic signal in the telomere lengths (Fig 2). Between plants with differing growth forms (herbaceous vs. woody) and plants with different ploidy levels (diploid vs. polyploid), λ was similar and elevated at levels greater than 0.75. But when comparing plants with different life spans, annual plants had a lower λ (0.57, 95% CI 0.28 – 0.83) compared to perennial plants (0.91, 95% CI 0.84 – 0.95). In addition, monocots had lower λ (0 and 95% CI 0 – 0.52) compared to eudicot plants (0.79, 95% CI 0.66 – 0.88). This indicated annual plants and monocots had telomere lengths that were more different between closely related species on average compared to distant relatives.

**Figure 2.**
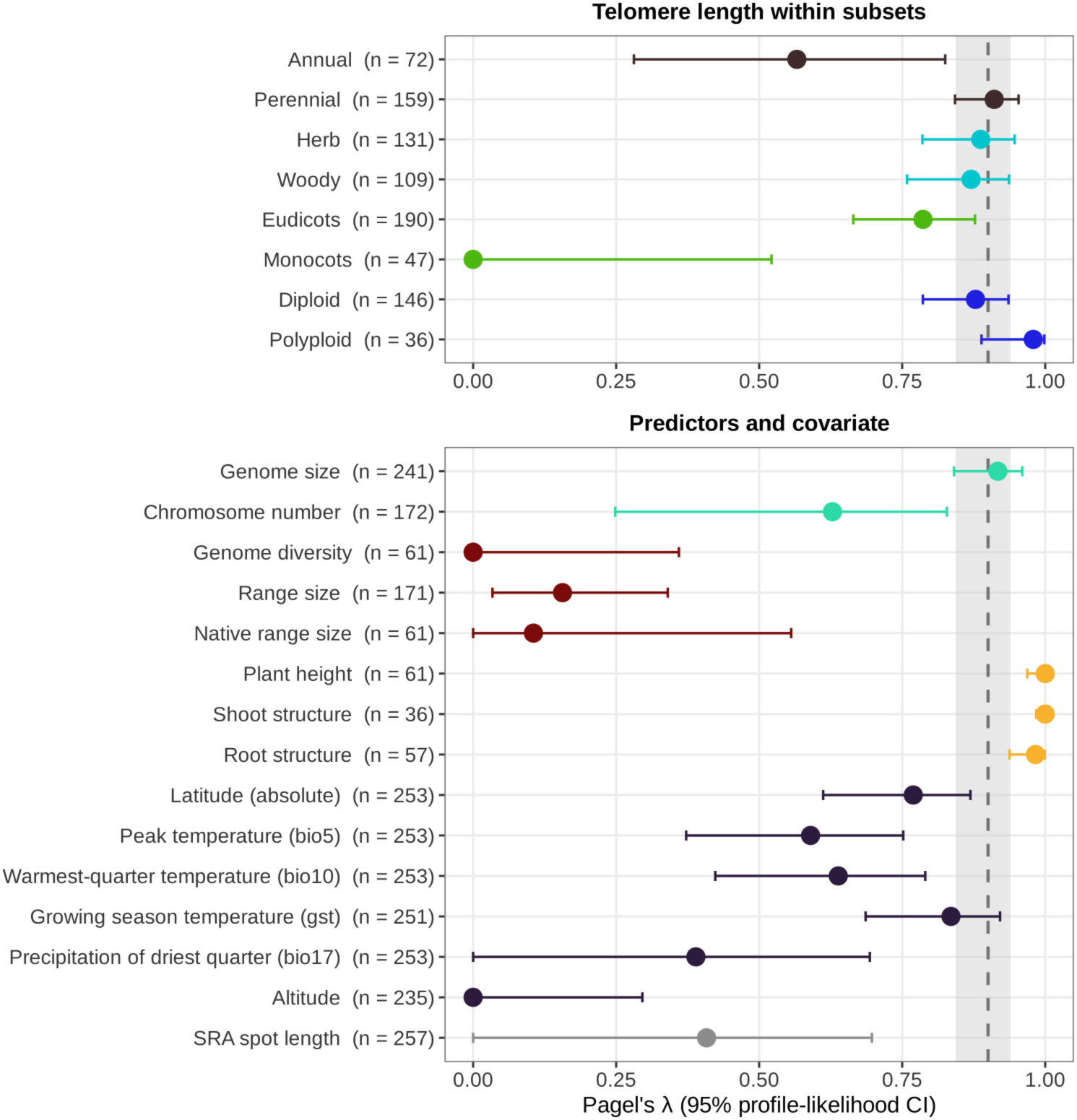
Distribution of phylogenetic signals (λ) for telomere length within life-history categories (top) and for different traits (bottom). λ value of 1 indicates the trait follows a Brownian motion model while λ of 0 indicates no phylogenetic structure in the traits. Different categories were separated using different colors. The dot represents λ estimated by maximum likelihood and bars represent the 95% confidence interval. Sample sizes are given beside each label. The dashed line and grey band show telomere length λ across all land plants (λ = 0.90, 95% CI 0.84 – 0.94). Latitude is absolute latitude, while altitude and native range size are log10-transformed.

We then estimated λ for other variables and compared against the telomere length λ to understand the macroevolutionary patterns spanning the plant kingdom (Fig 2). For the plant species we analyzed, we estimated λ for two molecular traits (*i.e.* genome size and chromosome number), three demography variables (*i.e.* genome diversity, range size and native range size), three physiological traits (*i.e.* height, shoot and root structure), six ecological parameters (*i.e.* peak temperature, warmest-quarter temperature, growing season temperature, precipitation of the driest quarter, latitude and altitude), and the technical covariate spot length (Fig 2 and Table S4). Results showed demography had the lowest λ values suggesting closely related species are not sharing the same demographic history and range sizes, which are consistent with previous findings (Webb and Gaston 2003). The ecological parameters had variable λ values, similar to observations from other plant studies (Liu et al. 2015; Koski and Ashman 2016; Steinbauer et al. 2016; Xu et al. 2019; Harris et al. 2022), where latitude (λ = 0.77) and growing season temperature (λ = 0.84) had high phylogenetic signal, peak and warmest-quarter temperature had intermediate signal (λ = 0.59 and 0.64), and precipitation of the driest quarter (λ = 0.39) and altitude (λ = 0) had low or no phylogenetic signal. The molecular and morphological traits had elevated λ consistent with prior studies as well (Kang et al. 2014; Li et al. 2017; Morton et al. 2024). Spot length had weak and non-significant phylogenetic signal (λ = 0.41, 95% CI 0 – 0.70).

### Ecological variables but not life-history were associated with telomere length

We examined if variation in plant life-history strategy was associated with differences in telomere length, and compared the telomere lengths between plant species with differing life span (median telomere length annual = 5,918.75 bp and perennial = 6,295 bp), growth form (median telomere length herb = 5,245 bp and woody = 6,706.25 bp), and ploidy (median telomere length diploid = 6,190 bp and polyploid = 5,236.25 bp). Growth form (Mann Whitney U test p-value = 7.8e-4) and ploidy (p-value = 0.023) differed in telomere length, but after accounting for phylogeny and spot length none of the life-history variables was associated with telomere length (PGLS lifespan p = 0.96, growth form p = 0.088, ploidy p = 0.77; Fig S2).

We further investigated the evolutionary forces that shaped telomere length in plants by regressing telomere length on 34 variables relating to the genome, life-history traits, range, or the ecological conditions of the plant. We used phylogenetic generalized least square (PGLS) regression to incorporate the underlying phylogenetic signal with the regression modeling (Revell 2010) with log10 spot length as a covariate to account for sequencing read length (Table 1; results without the covariate and OLS results are in Table S5). After multiple hypothesis testing correction for testing 34 variables (Benjamini–Hochberg), the mean daily maximum temperature of the warmest month (bio5) was significantly and positively associated with telomere length (PGLS adjusted p = 0.018), and the mean temperature of the warmest quarter (bio10) was positively associated at marginal significance (adjusted p = 0.050) (Fig 3A, B).

**Figure 3.**
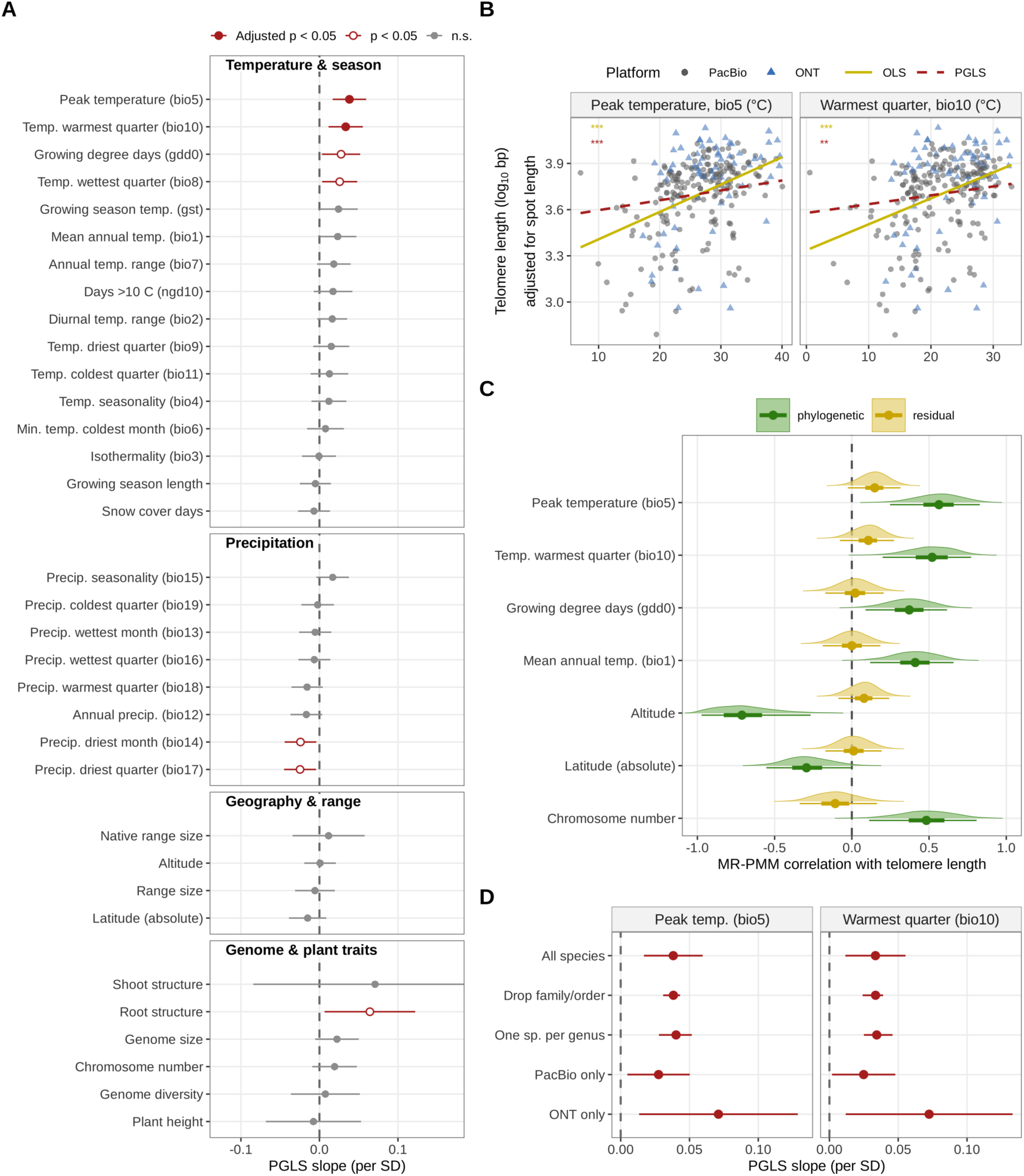
Association between telomere length and predictor variables. (A) PGLS slopes (change in log10 telomere length per SD of each predictor, 95% CI) for all 34 predictors with log10 spot length as a covariate. Filled red: significant after correction for the effective number of independent tests (Li and Ji 2005). Open red: p < 0.05 before correction. Grey: not significant. (B) Telomere length adjusted for spot length against peak temperature (bio5) and warmest-quarter temperature (bio10). Circles represent estimates from PacBio platform and triangles represent estimates from ONT platform. Solid gold line, OLS; dashed red line, PGLS (drawn through the data centroid with the fitted slopes). Asterisks give significance for OLS (gold) and PGLS (red): *** p < 0.001, ** p < 0.01. (C) Posterior distributions of the phylogenetic (green) and residual (gold) correlations between telomere length and each predictor from bivariate MR-PMM, with spot length as a fixed effect on telomere length. Points represent posterior medians. Thick and thin bars represent 50% and 95% credible intervals. (D) Robustness of the bio5 and bio10 slopes fitted with a PGLS using all species, and compared against PGLS omitting one family or order at a time (median and range of 45 analyses), subsampling one species per genus (median and 95% range of 200 subsamples), or using only PacBio (n = 188) or ONT (n = 65) species. Bars are 95% CIs unless stated.

**Table 1.** Phylogenetic generalized least square (PGLS) regression of telomere length on genome, geographic, and climatic variables with log10 spot length as a covariate. Slopes are per standard deviation of each predictor (SE in parentheses). Adjusted p-values are Benjamini–Hochberg corrected across all 34 predictors (Table S5), and values significant after correction are in bold. λ, Pagel’s λ of the residuals. Results for the remaining predictors without the covariate and from OLS are in Table S5.

| Variable | N | Slope per SD (SE) | p | Adjusted p | λ |
|---|---|---|---|---|---|
| Absolute latitude | 253 | -0.015 (0.012) | 0.22 | 0.39 | 0.91 |
| Median altitude (log10 m) | 235 | 0.001 (0.010) | 0.94 | 0.96 | 0.90 |
| Genome size (log10 Mb) | 241 | 0.022 (0.014) | 0.12 | 0.28 | 0.91 |
| Chromosome number (2n) | 172 | 0.019 (0.014) | 0.18 | 0.37 | 0.74 |
| Mean annual temperature (bio1) | 253 | 0.023 (0.012) | 0.055 | 0.21 | 0.90 |
| Mean diurnal temperature range (bio2) | 253 | 0.016 (0.010) | 0.095 | 0.28 | 0.91 |
| Isothermality (bio3) | 253 | -0.001 (0.011) | 0.96 | 0.96 | 0.91 |
| **Max. temperature of warmest month (bio5)** | 253 | **0.038 (0.011)** | **5.3e-4** | **0.018** | 0.90 |
| Min. temperature of coldest month (bio6) | 253 | 0.008 (0.012) | 0.52 | 0.72 | 0.91 |
| Mean temperature of wettest quarter (bio8) | 253 | 0.026 (0.011) | 0.024 | 0.16 | 0.91 |
| Mean temperature of driest quarter (bio9) | 253 | 0.015 (0.012) | 0.20 | 0.38 | 0.90 |
| Mean temperature of warmest quarter (bio10) | 253 | 0.033 (0.011) | 0.003 | 0.050 | 0.90 |
| Mean temperature of coldest quarter (bio11) | 253 | 0.013 (0.012) | 0.30 | 0.48 | 0.91 |
| Precipitation of driest month (bio14) | 253 | -0.024 (0.010) | 0.020 | 0.16 | 0.92 |
| Precipitation seasonality (bio15) | 253 | 0.017 (0.011) | 0.11 | 0.28 | 0.91 |
| Precipitation of driest quarter (bio17) | 253 | -0.025 (0.010) | 0.018 | 0.16 | 0.92 |
| Mean temperature of growing season (gst) | 251 | 0.024 (0.012) | 0.052 | 0.21 | 0.91 |
| Growing degree days >0 °C (gdd0) | 253 | 0.028 (0.012) | 0.027 | 0.16 | 0.90 |
| Growing degree days >10 °C (ngd10) | 248 | 0.017 (0.013) | 0.17 | 0.36 | 0.90 |
| Snow cover days (scd) | 253 | -0.007 (0.010) | 0.51 | 0.72 | 0.91 |

Many climatic variables were strongly correlated with each other (Fig S3A), so the 34 predictors represented approximately 17 effective independent tests (Li and Ji 2005). After correcting for this effective number of tests, bio5 remained significant (adjusted p = 0.009) and bio10 was marginally significant (adjusted p = 0.0497; Table S5). For bio5, each 10 °C increase corresponded to approximately 16% longer telomeres. Growing degree days (gdd0), the temperature of the wettest quarter (bio8), precipitation of the driest month (bio14) and driest quarter (bio17), and root structure were nominally associated with telomere length (p < 0.05) but were not significant after multiple hypothesis correction (Table 1).

We then tested the robustness of the temperature associations by refitting the PGLS models on subsets of the data. Both bio5 and bio10 remained significant in all 45 datasets after removing each family or order with three or more species, Rosaceae and Brassicaceae together, or the bryophytes and the lycophyte. When we generated 200 random subsamples retaining one species per genus, 98% (bio5) and 89% (bio10) had significant PGLS results (Fig 3D; Table S6). Because ONT species also occupied warmer sites than PacBio species (PGLS p = 0.019 for bio5) we repeated the analysis within each platform. The bio5 slope was positive and significant in both PacBio (p = 0.018) and ONT species (p = 0.019) (Fig 3D) and the slopes did not differ significantly between platforms (bio5 × platform interaction p = 0.10; Table S7).

Including platform as an additional covariate retained the bio5 (p = 0.002) and bio10 (p = 0.008) associations, although neither remained significant after multiple-testing correction (Table S7).

### Analyses support peak temperature as the most informative predictor

To identify which of the correlated climatic variables best explained telomere length, we compared PGLS models that contained one climatic or geographic predictor plus spot length as a covariate and fitted to the 230 species with data for all predictors. The bio5 model was best supported (Akaike weight 0.81) followed by bio10 (ΔAICc = 4.0), whereas models with latitude or altitude fitted no better than a model with spot length alone (Fig S3B; Table S8). The temperature variables were strongly correlated with each other (Spearman ρ = 0.95 between bio5 and bio10, and 0.82 between bio5 and bio1) and moderately with latitude and altitude (Fig S3A). Accounting for multicollinearity, bio5, absolute latitude, and altitude were included together in a single PGLS model with spot length (n = 234), bio5 remained significant (p = 5.2e-4), whereas latitude (p = 0.66) and altitude (p = 0.49) did not. Bio5 remained significant when sequencing platform was also included (p = 0.002; Table S9).

To test whether bio5 reflected general warmth, winter cold, or temperature seasonality, we added mean annual temperature (bio1), the minimum temperature of the coldest month (bio6) or the annual temperature range (bio7) to the bio5 model. Bio5 remained significant in each model (p ≤ 0.002), whereas none of the added variables was significant (p ≥ 0.20; Table S9). Within clades, the bio5 slope was significant in clades rosids, fabids, and malvids; and orders Fabales, and Brassicales. The bio5 slope was positive but not significant in monocots, order Poales, and clade asterids, and close to zero in the orders Rosales and Malvales (Fig S4). The temperature association was also stronger in herbaceous species (27% longer telomeres per 10 °C increase in bio5, p = 2.6e-4) than in woody species (7%, p = 0.24), and in annual (33%, p = 0.002) than in perennial species (15%, p = 0.003), although the temperature × growth form (p = 0.069) and temperature × lifespan (p = 0.091) interactions were not significant (Table S10).

### Covariation between telomere length and temperature is phylogenetically structured

The regression analysis indicated temperature was a significant factor explaining telomere length variation in plants, whereas latitude was not associated with telomere length in the PGLS analysis. Because latitude had a strong phylogenetic signal in the plant species we analyzed (Fig 2), we hypothesized the shared phylogenetic signal with telomere length could have been a confounding factor during the PGLS analysis and mask covariation that is structured by phylogeny (Westoby et al. 2023). To investigate the phylogenetic conservation between traits (Losos 2008) we used a Bayesian multi-response phylogenetic mixed model (MR-PMM), which treat traits as joint response variables and decompose the trait correlations into phylogenetic and non-phylogenetic components (Halliwell et al. 2025). An elevated phylogenetic component indicates high covariation between traits were driven by shared phylogenetic history and shared selective pressure (Westoby et al. 2023).

We fitted bivariate MR-PMMs with telomere length and each predictor as joint responses using all species with absolute latitude and including log10 spot length as a fixed effect on telomere length (Fig 3C, Fig S5, and Table S11). Peak temperature had a significant positive phylogenetic correlation with telomere length (0.56 and 95% credible interval [0.26 – 0.84]), as did warmest-quarter temperature (0.52 [0.24 – 0.81]), mean annual temperature (0.41 [0.12 – 0.66]), growing degree days (0.37 [0.09 – 0.61]), the temperature of the wettest quarter (0.33 [0.04 – 0.64]) and growing season temperature (0.29 [0.03 – 0.57]). On the other hand, the corresponding residual (non-phylogenetic) correlations were close to zero. Latitude had a negative phylogenetic correlation with telomere length (−0.29 [-0.56 – −0.005]) but this estimate was sensitive to removing individual families (the 95% CI excluded zero in 8 of 15 reduced datasets), whereas the phylogenetic correlations of peak and warmest-quarter temperature excluded zero in 14 of 15 (Table S12). Altitude had a strong negative phylogenetic correlation (−0.71 [−1.00 – −0.34]), but only about 20% of the variance in altitude was phylogenetic and mixing was poor. Precipitation of the driest quarter had a significant negative residual correlation with telomere length (−0.21 [−0.38 – −0.03]), but neither precipitation variable had a significant phylogenetic correlation.

Previous studies have found genome size in plants is shaped by ecological differences (Greilhuber and Leitch 2013; Cacho et al. 2021; Bureš et al. 2024). Genome size and chromosome number were not associated with telomere length in the PGLS analysis (Table 1). We further investigated whether the size of the genome has a phylogenetic conservation with telomere length by conducting MR-PMM analysis. Results showed (Fig S5) only the chromosome number had a significant positive phylogenetic correlation with telomere length (0.48 and 95% CI [0.13 – 0.81]), whereas genome size did not (0.20, 95% CI [-0.06 – 0.46]).

### The temperature association is not driven by ancient divergences

As telomere length and bio5 were significantly divergent from Brownian model (λ = 0.9), we examined if the telomere length and temperature evolution were driven by evolutionary events deep in the phylogeny (Uyeda et al. 2018). An early-burst model where evolutionary rates decline through time did not explain telomere length variation (Table S3). Disparity through time analysis (Harmon et al. 2003) showed no early-burst signature for either trait, where the morphological disparity index (MDI) did not significantly differ from Brownian motion expectation (telomere length MDI = −0.19, p = 0.30; bio5 MDI = 0.21, p = 0.71). But approximately within the last 60 (telomere length) and 100 (bio5) million years the relative disparity within clades exceeded the Brownian motion expectation (Fig 4A). Node height tests (Freckleton and Harvey 2006; Slater and Pennell 2014) showed the magnitude of phylogenetically independent contrasts (Felsenstein 1985) increased toward the present (p = 3.3e-5 and p = 1.8e-12 respectively; Fig S6A and B). In addition, we regressed telomere length contrasts on bio5 contrasts (Garland et al. 1992) using a tree transformed by the PGLS λ separately for nodes younger and older than the median node age (25 million years). The association was significant for younger (126 nodes; slope = 0.0056, p = 0.016) and older divergences (126 nodes; slope = 0.0079, p = 0.0096; Fig 4B; Table S13). On the λ-transformed tree, telomere length contrasts were unrelated to node height (p = 0.53; Fig S6C). The association was also apparent among family means (Spearman ρ = 0.46, p = 0.031; Fig 4C) where the temperate families Rosaceae and Brassicaceae had the shortest telomeres, and families from warmer regions had longer telomeres across the major angiosperm clades. Much of the divergence in telomere length and temperature occurred among recently diverged lineages, in line with the phylogenetic signal of telomere length being below that expected under Brownian motion (λ = 0.90).

**Figure 4.**
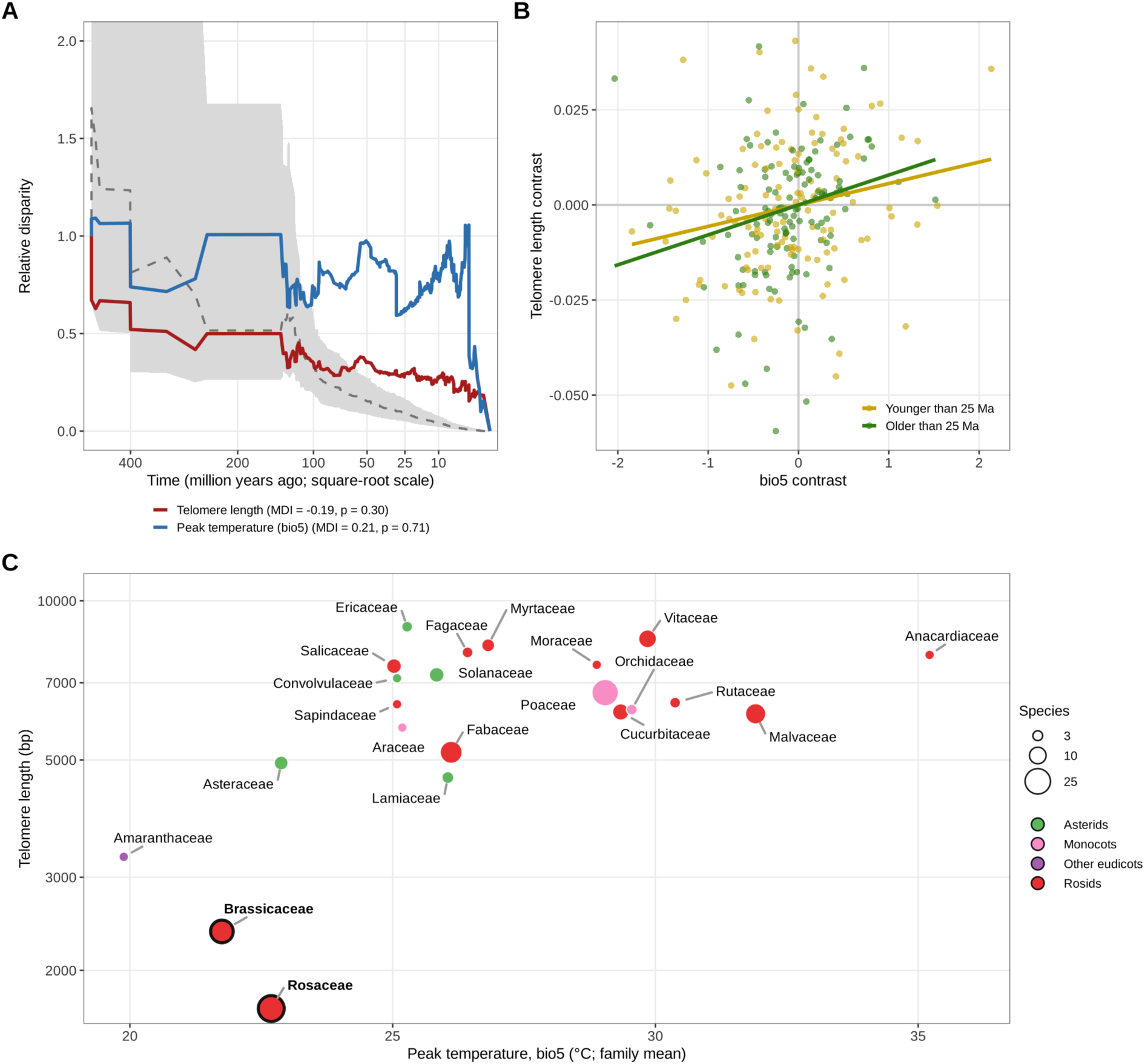
The temperature association through evolutionary time. (A) Disparity through time for telomere length (adjusted for spot length; red) and peak temperature (bio5; blue) on the time-calibrated tree (Harmon et al. 2003). Relative disparity is the mean disparity within clades present at each time divided by total disparity. Dashed line, median of 10,000 Brownian motion simulations; grey band, 95% range of the simulations. MDI, morphological disparity index with two-sided p-value. (B) Telomere length contrasts against bio5 contrasts computed on the tree transformed by the PGLS λ (0.90) for nodes younger (gold) and older (green) than the median node age (25 Ma). Lines, regressions through the origin. (C) Family means of telomere length (adjusted for spot length) against family mean peak temperature for families with ≥3 species.Point size, number of species; color, major clade; Rosaceae and Brassicaceae are bolded and outlined.

### Idiosyncratic effects of domestication on telomere length

Domestication has played a crucial role during the evolution of various species in both animal and plant kingdom (Purugganan 2022). Prior studies have suggested telomeres have lengthened in domesticated species (Pepke and Eisenberg 2022; Colt et al. 2024), suggesting a life-history relating to a domestication selection pressure have universally selected for increasing the length of the telomere. However, these studies have not compared domesticates to their direct wild progenitors and tested the effects of domestication on telomere length evolution. In yeast, wild isolates have significantly shorter telomeres compared to domesticated strains (D’Angiolo et al. 2022), suggesting domestication could have an evolutionary effect of lengthening telomeres.

We investigated the relationship between domestication and telomere length by comparing domesticated crops with their wild progenitors using population genomic sequencing data (Table S14). Because ONT libraries produce longer telomere length estimates than PacBio libraries we restricted the analysis to comparisons in which the domesticate and its wild progenitor were sequenced on the same platform and had the same ploidy with at least four genotypes per group. This retained six comparisons across five crops (Fig 5). Telomere length differed between domesticates and wild progenitors in only two comparisons and in opposite directions. Cultivated cotton (*Gossypium herbaceum*) had longer telomeres than wild *G. herbaceum* (median 13,873 vs. 7,013 bp; Mann–Whitney U test p = 0.009), whereas foxtail millet (*Setaria italica*) had shorter telomeres than its progenitor *S. viridis* (5,753 vs. 6,085 bp; p = 0.028). Tomato (*Solanum lycopersicum*), proso millet (*Panicum miliaceum*), and indica and japonica rice (*Oryza sativa*) did not differ significantly from their wild progenitors.

**Figure 5.**
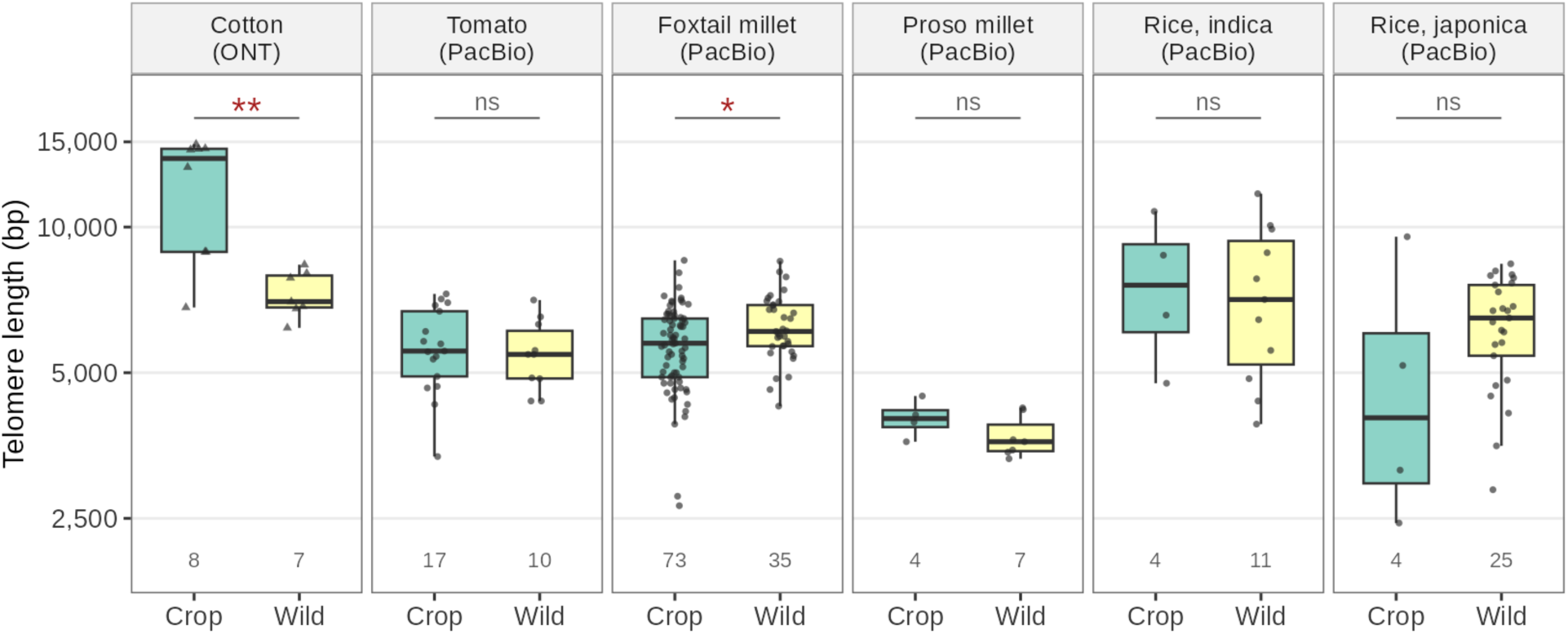
Telomere length of domesticated crops and their wild progenitors. Progenitors for each domesticates correspond to wild *Gossypium herbaceum* for cotton, *Solanum pimpinellifolium* for tomato, *Setaria viridis* for foxtail millet, wild *Panicum miliaceum* for proso millet, *Oryza rufipogon* Or-Ia for indica rice, and Or-IIIa for japonica rice. Points are individual genotypes and numbers give sample sizes. Asterisks denote Mann–Whitney U test significance (* p < 0.05, ** p < 0.01; ns, not significant). Triangles, ONT; circles, PacBio. The y-axis is log-scaled.

## Discussion

In this study we investigated the evolutionary force that shaped telomere length in plants by examining the length variation in species across the green plant kingdom. We discovered life-history variation between plants had minimal association with telomere length, whereas temperature was associated with global plant telomere length variation. Peak temperature in particular had a significant positive association with telomere length after accounting for phylogeny, sequencing read length, and multiple testing. It was a better predictor than latitude, mean annual temperature, or other climatic variables, pointing to peak summer heat rather than geography or winter cold as the relevant factor. These results suggested abiotic factors have a significant role in shaping telomere length in plants. The covariation between temperature and telomere length was largely phylogenetic in the MR-PMM analysis, which models phylogeny and trait evolution jointly (Westoby et al. 2023). The results indicated phylogenetic conservatism (*i.e.* related species share similar traits through common ancestry) (Losos 2008) and shared selective pressure through ecology (*i.e.* telomere length is selected along a temperature gradient) are acting together to shape the telomere length in plants. Importantly, the association was not concentrated at a few ancient divergences but was present among both recently and anciently diverged lineages, consistent with a repeated association between temperature and telomere length rather than a single historical event.

Latitude had a weaker and less robust negative association with telomere length and depended on temperate families (*i.e.* Brassicaceae and Rosaceae) with short telomeres. A latitudinal cline consistent with the temperature association was also observed within *Arabidopsis thaliana* populations where ecotypes closer to the equator and experience warmer growing seasons had longer telomeres compared to ecotypes closer to the Arctic circle (Choi et al. 2021). Altitude was not associated with telomere length in the PGLS analysis and its negative phylogenetic correlation in the MR-PMM should be interpreted cautiously because altitude carried little phylogenetic signal. Although its direction is consistent with shorter telomeres at colder and higher sites. Interestingly, genome size variation in plants is also associated with latitude (Levin and Funderburg 1979; Knight et al. 2005; Bureš et al. 2024). We found no significant association between telomere length and genome size in either the PGLS or MR-PMM analysis, suggesting a weak link between the two molecular traits, although chromosome number had a positive phylogenetic correlation with telomere length. Nevertheless, variation in telomere length and genome size that tracks climate suggests that environmental gradients can shape the evolution of fundamental properties of the chromosome.

Genome size variation in plants is significantly associated with several ecological factors, especially those relating to temperature (Bureš et al. 2024). For telomere length, we also found temperature to be significantly associated with telomere length after correcting for phylogenetic relatedness. Pine trees were found to have a similar trend between telomere length and mean temperature of different seasons (Mu et al. 2015). Based on our findings we propose the ecological adaptation model of plant telomere length, where we posit selective forces arising from abiotic factors drives the variation of telomere length in plants. Temperature, in particular, could be an important ecological factor that drives the global plant telomere length.

Our telomere length results contrast the findings from animal telomere length studies that have found significant associations with life-history traits (Gomes et al. 2011; Pepke and Eisenberg 2022). We initially predicted differences in life-history strategies would also have significant effects on telomere length variation in plants, for example, differences in meristem importance for annual and perennial plants with contrasting reproductive modes could lead to differential selection and telomere length differences between growth types. But in plants, we found no significant associations between differences in growth strategy, demography, or phenotypic traits with telomere length. This might be explained if selection on the meristem is universal across plants regardless of life-history strategy. Recent studies have suggested most plants may have a segregated germline (Lanfear 2018) and the protection of meristematic stem cells through the telomere, if existing, might be universal with no selective differences between plants with differing growth forms. The life-history variables available for most species were coarse (*e.g.* annual vs. perennial) and continuous traits such as height and root or shoot structure were available for fewer than 65 species, so modest life-history effects cannot be excluded.

In the end, life-history variation may have a minor role in shaping telomere length variation in plants. Our domestication results also support this conclusion. Domestication has rapidly and drastically altered plant growth and life history, telomere length showed no consistent change between crops and their wild progenitors. On the other hand, the significant association between telomere length and flowering time in example plant species (Choi et al. 2021; Campitelli et al. 2022), might seem to contradict our findings that life-history difference does not explain telomere length differences in plants. But flowering time variation and growth differences in plants arose as an adaptation to the surrounding environmental conditions (Blackman 2017; Gaudinier and Blackman 2020). We suggest abiotic factors relating to biogeographic clines have a stronger selective force compared to the selection arising from life-history differences in shaping the telomere length variation in plants.

The biological mechanism that links temperature with telomere length is unknown and a natural focus for future research. Hints, however, can be gained from the association between temperature and global telomere length variation where plants from cooler climates have shorter telomeres on average. Plants in low temperature have slower cellular division and longer generation time (Bennett et al. 1982; Francis and Barlow 1988). Consistent with this idea, the temperature association was about three times steeper in herbaceous than in woody species and steeper in annual than in perennial species, although these differences were not statistically significant (Fig S4). If changes in the telomere length underlies the changes in cellular development under varying temperature, this could be the selective force that shapes telomere length variation in plants. It is unclear if the length of the telomere itself confers selective benefit or if selection is occurring at the telomerase activity (Kumawat and Choi 2023). In the latter selection scenario, if a specific ecological condition selects for higher (or lower) telomerase activity, an increase (or decrease) in telomere length is a consequence of the telomerase activity and not the direct target of selection. A future direction that investigates the telomerase activity under various ecological conditions of different plant species would result in fruitful mechanistic insights underlying the ecological adaptation model of plant telomeres.

## Methods

### Estimating telomere length across the plant kingdom

We searched for plant species with long read sequencing data by searching the N3 database (Xie et al. 2024). The telomere sequence in plants are highly variable (Shakirov et al. 2022) and to avoid any biases that could arise from telomere sequence differences we focused on plant species with the ancestral *A. thaliana* type telomere repeat sequence TTTAGGG (Richards and Ausubel 1988). Telomere sequence of a plant was searched at TeloBase (Lyčka et al. 2024). Using the long read sequencing data telomere length of each plant species was estimated using Topsicle (Nguyen and Choi 2025). We use the parameter telomere repeat count (TRC) values higher than 0.7 (--cutoff 0.7) as a cutoff for deciding telomere originating sequencing reads. In our previous study a TRC cutoff of 0.7 was deemed a conservative cutoff for plant and animal long read sequences (Nguyen and Choi 2025). The 5-mers of the telomere repeat sequence TTTAGGG were used to accommodate potential error or sequence variation within the telomere region. Sequencing reads with a minimum length of 7 kbp (--minSeqLength 7000) and maximum possible telomere length of 30 kbp (--maxlengthtelo 30000) were used for estimating telomere length. For the Charophyta and Chlorophyta orders previous study suggested species from these orders might have telomeres shorter than 1000 bp (Chaux et al. 2023), and to accommodate this shorter length we used TRC of 0.4 to relax the threshold and have the minimum length of sequencing reads for analysis as 3 kbp. Species with sequencing libraries that had at least 50 sequences that were classified as telomere containing long reads were used for estimating the telomere length. Every long reads classified as containing the telomere, we estimated the telomere length using the binary segmentation algorithm implemented in Topsicle. For a given species its telomere length was calculated as the median telomere length estimates from all telomere long reads identified and analyzed by Topsicle.

### Phylogeny reconstruction and estimating the phylogenetic signal

Species nomenclature considered in this study follows the World Plant database (https://www.worldplants.de). Species were assigned to clades by order following APG IV: monocots; rosids (fabids, malvids and Vitales); fabids (Rosales, Fabales, Malpighiales, Cucurbitales and Fagales); malvids (Brassicales, Malvales, Sapindales and Myrtales); and asterids (Lamiales, Solanales, Ericales, Asterales, Apiales, Boraginales, Cornales, Dipsacales and Garryales). Orders with at least 15 species (Poales, Rosales, Fabales, Brassicales and Malvales) were also analyzed separately.

We reconstructed the phylogenetic tree of the species estimated for their telomere length using time-calibrated supertrees (Zanne et al. 2014; Smith and Brown 2018). The supertree was pruned using the V.PhyloMaker2 package in R (Jin and Qian 2022) with the GBOTB.extended.WP option. Since V.PhyloMaker2 was built for vascular plants, phylogenetic tree and divergence time for non-vascular land plants were reconstructed based on previous studies (Heinrichs et al. 2007; Villarreal A. et al. 2016; Morris et al. 2018). With the phylogenetic tree we used the phytools package v2.4-4 in R (Revell 2012) to examine the phylogenetic signals of telomere length and other life history traits of the species included in this study.

Pagel’s lambda (Pagel 1997) was calculated using a maximum-likelihood approach implemented in phytools. Confidence intervals for λ were obtained from the profile likelihood. For telomere length we searched for the best fitting evolutionary model using the fitContinuous function from geiger package in R (Pennell et al. 2014). The best fitting model was decided as the one with the lowest corrected Akaike information criterion (AICc).

### Life-history and biogeography variables

Species occurrences were recorded by initially searching for the latitude and longitude coordinates from the original study that published the genome sequencing data. If there was no information from the published study species occurrence were derived from the Global Biodiversity Information Facility (GBIF) database using rgbif v3.8.5 (Chamberlain et al. 2025). The representative occurrence or hotspot of species occurrence was determined by 2D kernel density estimation on all filtered occurrences with grid size of 100 using the kde2d function in MASS package in R. Altitude was estimated as median elevation from the species occurrence data. Absolute latitude was used in all comparative analyses and altitude and native range size were log10-transformed.

Biogeographic parameters were obtained from the CHELSA-bioclim database v2.1 (Brun et al. 2022) based on the latitude and longitude coordinate for each species using the terra package in R (Hijmans et al. 2026). Plant clade, order, and family information were extracted from the N3 database (Xie et al. 2024), contig N50 statistics of the assembled genome and genome size were extracted from N3 database (Xie et al. 2024), chromosome number were extracted from (Bureš et al. 2024), root and shoot statistics were extracted from (Tumber-Dávila et al. 2022).

Life span and growth form were extracted from several sources (Tumber-Dávila et al. 2022; Poppenwimer et al. 2023; Bureš et al. 2024) and demographic parameters were extracted from (Roberts and Josephs 2025).

### Statistical analysis

All statistical analyses and visualizations were done in R (v4.3.3). Ordinary least square analyses were done using the lm function. Mann–Whitney U tests were conducted in R. PGLS models estimated Pagel’s λ for the residuals by maximum likelihood using phylolm (Tung Ho and Ané 2014). Predictors were standardized to unit variance so that slopes are comparable among variables. To account for technical variation, all comparative models included log10 of the SRA library average spot length (median across the runs analysed for each species; Table S15) as a covariate. For analyses of single traits, telomere length was adjusted for spot length by subtracting the spot length term of the PGLS model. Sequencing platform was coded as ONT (GridION, MinION, PromethION) or PacBio (RS, RS II, Sequel, Sequel II, Revio). P-values across the 34 predictors were adjusted with the Benjamini–Hochberg procedure.

To account for correlation among predictors, we estimated the effective number of independent tests from the eigenvalues of the Spearman correlation matrix of the 34 predictors (Meff = 17.2) (Li and Ji 2005) and adjusted using Dunn-Šidák correction (Table S5). Robustness was assessed by refitting PGLS models after removing each family or order with at least three species, Rosaceae and Brassicaceae together, and the bryophytes plus lycophyte (45 reduced datasets), and on 200 random subsamples retaining one species per genus.

The Bayesian MR-PMM analysis was conducted using the MCMCglmm R package (Hadfield 2010) with the Gaussian multivariate normal distribution and parameters were set at 1,100,000 iterations with burn-in of 100,000 and thinning of 1,000. The phylogenetic covariance used a parameter-expanded prior and the residual covariance an inverse-Wishart prior (V = I, ν = 2). The log10 spot length was included as a fixed effect on telomere length. Each model was run as two independent chains. Sensitivity to individual clades was assessed by refitting each model after removing each family with at least five species, Rosaceae and Brassicaceae together, and the bryophytes plus lycophyte (Table S12). Since Bayesian analyses do not return significance statistics MR-PMM bivariate models were validated by visually inspecting MCMC chains of posterior estimates and confirming that the effective sample size of the correlation estimates exceeded 200 in every model (Table S12). The posterior median trait distribution, the 50% and 95% credible interval (CI) were derived from the model. The estimates were deemed significant when 95% CI does not contain point 0.

### Disparity through time, node height tests, and contrast regressions

To test whether the temperature association was driven by ancient divergences we analyzed telomere length (adjusted with spot length) and bio5 with three approaches and computed phylogenetically independent contrasts (Felsenstein 1985) using ape (Paradis and Schliep 2019). Disparity through time was computed with the dtt function of geiger (Harmon et al. 2003; Pennell et al. 2014) on the time-calibrated tree using the average squared Euclidean distance among species as the disparity index. As estimating morphological disparity index (MDI) at various time points is prone to high false-positive rate (Type 1 error), rank envelope method was performed with 10,000 Brownian motion simulations to achieve confidence range (Murrell 2018). The area between the observed curve and confidence range was compared using a two-sided test. Node height test (Freckleton and Harvey 2006) regressed the log absolute standardized contrasts of each trait on node height above the root with ordinary and robust regression using the nh.test function of geiger (Pennell et al. 2014; Slater and Pennell 2014). Contrasts that had a value of zero (three for bio5) were set to a minimal value of 10⁻⁵ for nh.test. Node height tests were run on the time calibrated tree for telomere length and bio5 and on the λ-transformed tree for telomere length. The association between the two traits was estimated by regressing telomere length contrasts on bio5 contrasts through the origin (Garland et al. 1992) using contrasts from the λ-transformed tree (Garland and Ives 2000; Blomberg et al. 2012). This regression was fitted separately to nodes younger and older than the median node age (25 Ma).

### Predictor comparison and within-group analyses

Collinearity among predictors was assessed with Spearman correlations. Climatic and geographic predictors were compared by fitting PGLS models containing each predictor and log10 spot length to the 230 species with complete data for all of them using AICc and Akaike weights. Multi-predictor PGLS models included bio5 together with absolute latitude and altitude, or together with one other temperature variable. Collinearity was assessed with variance inflation factors. Slopes within clades (monocots, Poales, rosids, fabids, malvids, asterids and orders with at least 15 species) and within life-history groups were estimated by refitting the bio5 and bio10 models with spot length in each group. Temperature × growth form and temperature × lifespan interactions were tested in PGLS models that included spot length.

### Domestication comparisons

Telomere lengths of domesticated crops and their wild progenitors were estimated with Topsicle from population genomic long-read sequencing data (Table S14). Because telomere length estimates differ between ONT and PacBio libraries, comparisons were restricted to domesticates and wild progenitors sequenced on the same platform and of the same ploidy with at least four genotypes per group. Telomere lengths of domesticates and wild progenitors were compared with two-sided Mann–Whitney U tests.

## Data availability

SRA ID of raw long read sequences analyzed in this study were reported in Table S1. Species telomere length estimate along with record of life-history traits, occurrence and bioclimatic variables are reported in Table S15. Analysis scripts are available from https://github.com/jaeyoungchoilab/PlantTeloLenEvo.git

## Acknowledgements

We thank members of the Choi lab Naseem Samo, Askhan Shametov, and Vandana Gurung for helpful suggestions during this study. This work was supported by a grant from the National Institute of General Medical Sciences of the National Institutes of Health R35GM154595 to J.Y.C.

---

**Figure S1. Sequencing read length and platform affect telomere length estimates.** (A) Telomere length against the median SRA spot length of each species’ libraries. Triangles, ONT; circles, PacBio. Solid gold line, OLS; dashed red line, PGLS. (B) Telomere length of species sequenced with ONT and PacBio.

**Figure S2. Telomere length across growth form, lifespan, and ploidy categories.** Boxes show the median and interquartile range. Significance is shown for Mann–Whitney U tests and for PGLS with log10 spot length as a covariate (*** p < 0.001, * p < 0.05; ns, not significant). The y-axis is log-scaled.

**Figure S3. Collinearity among predictors and comparison of climatic and geographic predictors.** (A) Spearman correlations among the 34 predictors ordered by hierarchical clustering. The predictors bio5 and bio10 are indicated in red. (B) ΔAICc of PGLS models containing each climatic or geographic predictor with log10 spot length as covariate fitted to a common set of 230 species relative to the best model (bio5). Filled red, predictor p < 0.05; open, not significant; black square and dashed line, model with spot length only.

**Figure S4. Temperature slopes within clades and life-history groups.** PGLS slopes (log10 telomere length per °C, ±95% CI) of telomere length on peak temperature (bio5) and warmest-quarter temperature (bio10) with log10 spot length as a covariate. Filled, p < 0.05; open, not significant; dotted line, all-species slope. Poales is nested in monocots; fabids and malvids in rosids; Rosales and Fabales in fabids; Brassicales and Malvales in malvids.

**Figure S5. Posterior distributions of the phylogenetic (green) and residual (gold) correlations.** MR-PMM model fit between telomere length and each of the 13 predictors with log10 spot length as a fixed effect on telomere length. Points, posterior medians; thick and thin bars, 50% and 95% credible intervals.

**Figure S6. Node-height tests.** Log absolute standardized contrasts of telomere length (A) and bio5 (B) against node age on the time-calibrated tree. Red lines, robust regression. (C) Node-height test for telomere length on the tree transformed by the PGLS λ (0.90).

**Table S1. Long-read sequencing libraries analyzed.** One row per SRA library with species, SRA accession, file size, number of reads and bases, and average spot (read) length.

**Table S2. Associations of telomere length estimates with sequencing variables.** PGLS regressions of log10 telomere length on log10 SRA spot length and sequencing platform (ONT vs PacBio; the estimate is the ONT effect in log10 units), and the Spearman correlation (ρ) between telomere length and contig N50 of the genome assembly. λ, Pagel’s λ of the PGLS residuals.

**Table S3. Comparison of evolutionary models for telomere length.** Models were fitted to log10 telomere length of the 257 land plants with fitContinuous. Pagel’s λ, phylogenetic signal; Pagel’s κ, branch length scaling; Ornstein–Uhlenbeck, attraction to an optimum (α, strength of attraction); rate trend, linear change in evolutionary rate through time; mean trend, directional drift; early burst, exponential decline in evolutionary rate through time; white noise, no phylogenetic covariance. The best-fitting model (lowest AICc) is in bold.

**Table S4. Phylogenetic signal (Pagel’s λ) of telomere length within life-history and clade subsets and of other traits.** λ was estimated by maximum likelihood with 95% profile-likelihood confidence intervals. The p-values are from likelihood-ratio tests against λ = 0. Latitude is absolute latitude with altitude and native range size are log10 transformed.

**Table S5. Regressions of log10 telomere length on each of the 34 predictors.** PGLS with log10 SRA spot length as a covariate (main analysis; Table 1), PGLS without covariates, and OLS without covariates. Predictors were standardized so slopes are the change in log10 telomere length per SD. p (BH), Benjamini–Hochberg adjusted p-value; p (Li & Ji), Šidák adjusted for the effective number of independent tests (Meff = 17.2; Li and Ji 2005). λ, Pagel’s λ of the residuals.

**Table S6. Robustness of the PGLS associations to removing clades and genus-level subsampling.** All models include log10 spot length as a covariate and slopes are per SD of each predictor. Reduced data sets were formed by removing each family or order with ≥3 species, Rosaceae and Brassicaceae together, and the bryophytes plus lycophyte together (45 data sets). Subsampling retained one randomly chosen species per genus (200 subsamples).

**Table S7. Robustness of the PGLS associations to sequencing platform.** PGLS with log10 spot length as a covariate fitted separately to PacBio and ONT species (slopes per SD of the predictor in the full data set), the p-value of the predictor × platform interaction, and PGLS with both log10 spot length and platform as covariates. p (BH), Benjamini–Hochberg adjusted p-values.

**Table S8. AICc comparison of single climatic and geographic predictors.** Comparing PGLS model of log10 telomere length containing one predictor and log10 spot length fitted to the 230 species with data for all predictors versus a model with spot length only. ΔAICc is relative to the best model. Akaike weights are across all models.

**Table S9. Multi-predictor PGLS models of log10 telomere length.** Each model includes log10 spot length. Predictors were standardized (slopes per SD). Platform is coded as ONT vs PacBio. VIF, variance inflation factor.

**Table S10. Temperature slopes within clades and life-history groups.** PGLS of log10 telomere length on peak temperature (bio5) or warmest-quarter temperature (bio10) with log10 spot length as a covariate fitted within each group. The % change in telomere length per 10 °C is 100 × (10^(10 × slope) − 1). The interaction p-value is for temperature × growth form (herb and woody rows) or temperature × lifespan (annual and perennial rows) in a model fitted to all species. Poales is nested in monocots; fabids and malvids in rosids; Rosales and Fabales in fabids; Brassicales and Malvales in malvids.

**Table S11. Phylogenetic and residual correlations between telomere length and each predictor from bivariate MR-PMMs.** Posterior medians and 95% highest posterior density intervals from MCMCglmm models with log10 spot length as a fixed effect on telomere length (two chains of 1,100,000 iterations, burn-in 100,000, thinning 1000). Phylogenetic share, the proportion of the predictor’s variance attributed to phylogeny; ESS, effective sample size; R-hat, Gelman–Rubin diagnostic.

**Table S12. Phylogenetic correlations from MR-PMMs after removing individual clades.** Posterior medians and 95% highest posterior density intervals after removing each family with ≥5 species, Rosaceae and Brassicaceae together, or the bryophytes plus lycophyte together. The first row repeats the full-data estimates in Table S11. Each reduced dataset was fitted as chains of 1,100,000 iterations (burn-in 100,000, thinning 1,000) so that the effective sample size exceeded 200.

**Table S13. Regressions of telomere length contrasts on bio5 contrasts by node age.** Regressions through the origin of phylogenetically independent contrasts of spot length adjusted log10 telomere length on contrasts of bio5. Contrasts were computed on a tree transformed by the PGLS λ (0.90) for all nodes and for nodes younger or older than the median node age (25 million years). The all-node slope equals the PGLS slope.

**Table S14. Telomere length of domesticated and wild genotypes in the crop–progenitor comparisons.** One row per sequencing library in the six comparisons shown in Fig 5 with species, SRA accession, platform, ploidy and Topsicle telomere length estimate.

**Table S15. Telomere length, sequencing, life-history, geographic and bioclimatic data for the 264 species.** One row per species. Telomere length is the median Topsicle estimate across telomere containing reads and SRA spot length is the median average spot length across the runs analyzed (*i.e.* log10 technical covariate). Bioclimatic variables (bio1–bio19, gst, gsl, gdd0, ngd10, scd) are from CHELSA v2.1 at the representative occurrence of each species.

